# 3D *in situ* imaging of female reproductive tract reveals molecular signatures of fertilizing spermatozoa in mice

**DOI:** 10.1101/2020.08.14.251736

**Authors:** Lukas Ded, Jae Yeon Hwang, Kiyoshi Miki, Huanan F. Shi, Jean-Ju Chung

**Affiliations:** Department of Cellular & Molecular Physiology, Yale School of Medicine, New Haven, CT, USA; Department of Obstetrics, Gynecology, and Reproductive Sciences, Yale School of Medicine, New Haven, CT, USA; Laboratory of Reproductive Biology, Institute of Biotechnology CAS, v.v.i., BIOCEV, Vestec, Czech Republic; Boston Children’s Hospital, Boston, MA, USA

## Abstract

Out of millions of ejaculated sperm, only a few reach the fertilization site in mammals. Flagellar Ca^2+^ signaling nanodomains, organized by multi-subunit CatSper calcium channel complexes, are pivotal for sperm migration in the female tract, implicating CatSper-dependent mechanisms in sperm selection. Here, using biochemical and pharmacological studies, we demonstrate that CatSper1 is an O-linked glycosylated protein, undergoing capacitation-induced processing dependent on Ca^2+^ and phosphorylation cascades. CatSper1 processing correlates with protein tyrosine phosphorylation (pY) development in sperm cells capacitated *in vitro* and *in vivo*. Using 3D *in situ* molecular imaging and ANN-based automatic detection of sperm distributed along the cleared female tract, we demonstrate that all spermatozoa past the UTJ possess intact CatSper1 signals. Together, we reveal that fertilizing mouse spermatozoa *in situ* are characterized by intact CatSper channel, lack of pY, and reacted acrosomes. These findings provide molecular insight into sperm selection for successful fertilization in the female reproductive tract.

## Introduction

In most mammals, millions or billions of spermatozoa are deposited into the cervix upon coitus. Yet less than 100 spermatozoa are found at the fertilization site, called ampulla, and only 10-12 spermatozoa are observed around an oocyte (Kolle, 2015; Suarez, 2002). This implies the presence of mechanisms to select sperm as they travel through the female reproductive tract and to eliminate non-fertilizing, surplus spermatozoa once the egg is fertilized (Sakkas et al., 2015). Recent *ex vivo* imaging studies combined with mouse genetics have shown that surface molecules on the sperm plasma membranes such as ADAM family proteins are essential for the sperm to pass through the utero-tubal junction (UTJ) (Fujihara et al., 2018). By contrast, whether such selection and elimination within the oviduct requires specific molecular signatures and cellular signaling of spermatozoa is not fully understood.

Mammalian sperm undergo capacitation, a physiological process to obtain the ability to fertilize the egg, naturally inside the oviduct (Austin, 1951; Chang, 1951). The emulation of sperm capacitation *in vitro* led to the development of *in vitro* fertilization (IVF) techniques (Steptoe and Edwards, 1976; Wang and Sauer, 2006). Since then, most studies on sperm capacitation and gamete interaction have been carried out under *in vitro* conditions. However, mounting evidence suggests that *in vitro* sperm capacitation does not precisely reproduce the time- and space-dependent *in vivo* events in the oviduct. Protein tyrosine phosphorylation (pY), which has been utilized as a hallmark of sperm capacitation over decades, showed different patterns in boar sperm capacitated *in vitro* from *ex vivo* and *in vivo* (Luno et al., 2013). In mice, pY is not required for sperm hyperactivation or fertility (Alvau et al., 2016; Tateno et al., 2013). Previous *in vitro* studies that represent the population average at a given time may or may not have observed molecular details of a small number of the most fertilizing sperm cells.

Capacitation involves extensive sperm remodeling that triggers cellular signaling cascades. Cholesterol shedding and protein modifications occur within the plasma membrane (Visconti et al., 1999; Vyklicka and Lishko, 2020). Cleavage and/or degradation of intracellular proteins by individual proteases and ubiquitin-proteasome system (UPS) also participate in the capacitation process (Honda et al., 2002; Kerns et al., 2016). Various capacitation-associated cellular signaling pathways that include cAMP/PKA activation followed by pY increase and rise in intracellular pH and calcium result in physiological outcomes such as acrosome reaction and motility changes (Balbach et al., 2018; Puga Molina et al., 2018). The sperm-specific CatSper Ca^2+^ channel forms multi-linear nanodomains on the flagellar membrane, functioning as a signaling hub that links these events and motility regulation during capacitation (Chung et al., 2014). Sperm from mice lacking CatSper genes are unable to control pY development and fail to migrate past the UTJ (Chung et al., 2014; Ho et al., 2009). The presence and integrity of CatSper nanodomains, probed by CatSper1, correlate with sperm ability to develop hyperactivated motility (Chung et al., 2017; Chung et al., 2014; Hwang et al., 2019). It is not known how these molecular and functional events are coordinated in the individual sperm cells within the physiological context.

Here, we reveal that most fertilizing mouse spermatozoa *in situ* are molecularly and functionally characterized by an intact CatSper channel, lack of pY, and reacted acrosomes. Using biochemical and pharmacological analyses, we show that CatSper1 undergoes O-linked glycosylation during sperm differentiation and maturation. Capacitation induces CatSper1 cleavage and degradation dependent on Ca^2+^ influx and protein phosphorylation cascades. We find that CatSper1 processing correlates with pY development in the flagella among heterogenous sperm cells capacitated *in vitro* and *in vivo*. We use *ex vivo* imaging and microdissection to show that intact CatSper channel is indispensable for sperm to successfully reach the ampulla and for the acrosome to react. Finally, we use newly developed 3D *in situ* molecular imaging strategies and ANN approach to determine and quantify the molecular characteristics of sperm distributed along the female reproductive tract. We demonstrate that all spermatozoa past the UTJ are recognized by intact CatSper1 signals which are graded along the oviduct. These findings provide molecular insight into dynamic regulation of Ca^2+^ signaling in selection, maintenance of the fertilizing capacity, and elimination of sperm in the female reproductive tract.

## Results

### CatSper1 undergoes post-translational modifications during sperm development and maturation

We previously found that the CatSper channel complex is compartmentalized within flagellar membrane, creating linear Ca^2+^ signaling nanodomains along the sperm tail (Chung et al., 2017; Chung et al., 2014; Hwang et al., 2019). Caveolin-1, a scaffolding protein in cholesterol-rich microdomains, colocalizes with the CatSper channel complex but does not scaffold the nanodomain (Chung et al., 2014). The molecular weight and amount of CatSper1, but not the other CatSper subunits, declines during sperm capacitation (Chung et al., 2014; Figure 1B, D). To better understand the processing of CatSper1, we first examined CatSper1 protein expression in the testis and epididymis. Interestingly, the molecular weight of CatSper1 increases gradually during sperm development and epidydimal maturation (Figure 1A; *upper*), indicating that CatSper1 undergoes post-translational modifications. We next examined the nature of the modifications. Block of tyrosine phosphatases by sodium orthovanadate or addition of specific protein phosphatases, PP1 or PTP, does not change the molecular weight of CatSper1 (Figure1-figure supplement 1A). In contrast, when sperm membrane was subjected to enzymatic deglycosylation, O-glycosidase, but not PNGase F, shifts apparent molecular weight of CatSper1 to close to the CatSper1 band with the smallest molecular weight observed in testis (Figure 1A, B, Figure1-figure supplement 1B). These data suggest that CatSper1 in sperm is not a phosphoprotein but an O-linked glycosylated protein.

**Figure 1.**
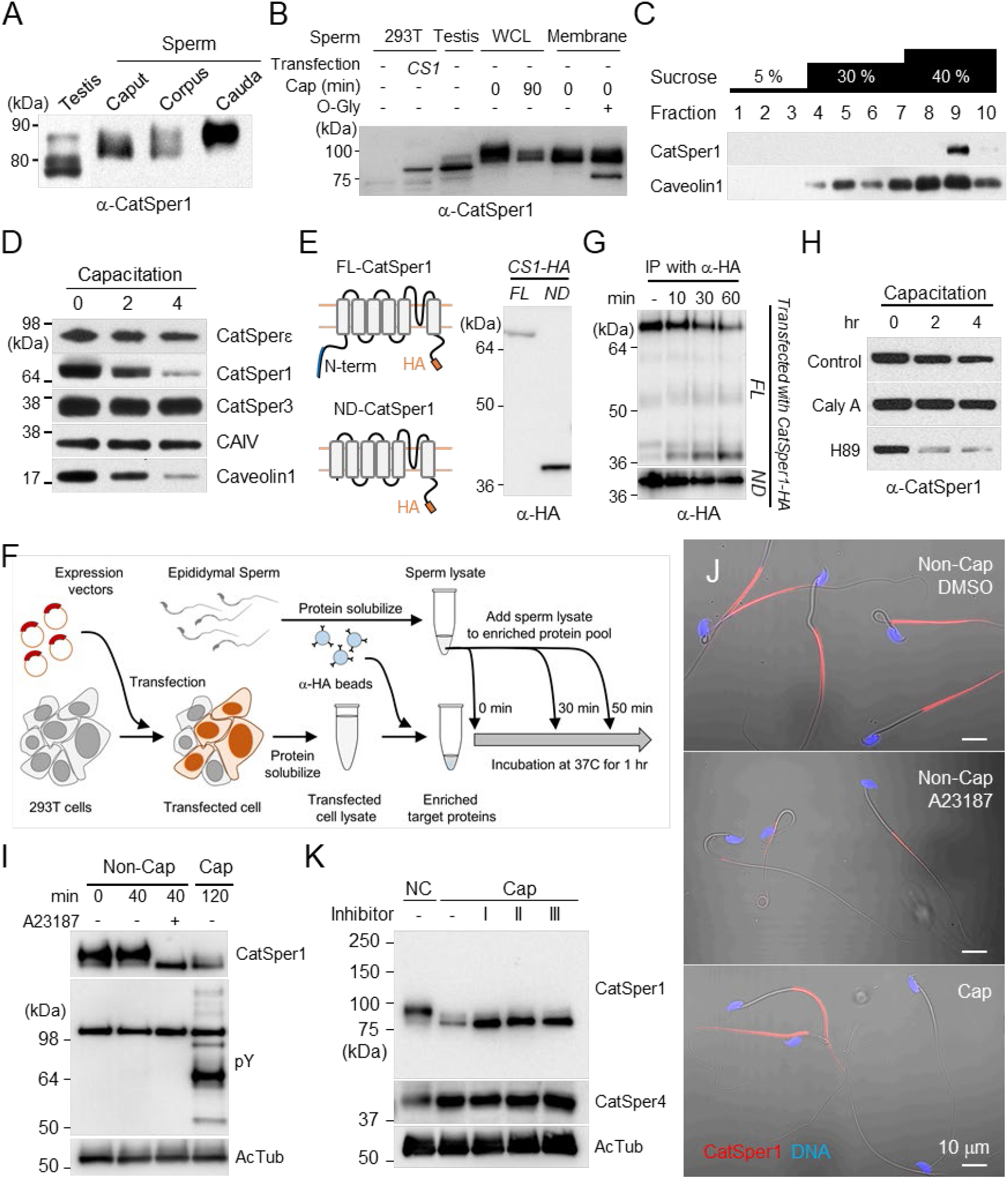
CatSper1 is specifically processed during *in vitro* capacitation. (A-B) CatSper1 undergoes post-translational modification during spermiogenesis and epididymal maturation. (A) A gradual decrease in electrophoretic mobility of CatSper1 is observed by Western blot analysis. (B) CatSper1 from sperm membrane fraction are shifted by O-glycosidase (O-Gly). (C) CatSper resides in lipid rafts subdomains of the plasma membrane in mature sperm. Solubilized sperm proteins were fractionized by discontinuous sucrose density gradient (5, 30, and 40%) centrifugation. (D) CatSper1 is degraded during the late stage of capacitation. Protein expression levels of CatSper1 and caveolin-1, but not CatSper3, CatSperε, or carbonic anhydrase 4 (CAIV) are altered by *in vitro* capacitation. (E-G) CatSper1 is cleaved within the N-terminal domain (NTD). (E) A cartoon of full-length (FL, *top*) and N-terminal truncated (ND, *bottom*) recombinant CatSper1 protein expressed in the study *(left*). Both proteins are tagged with HA at their respective C-termini (orange). CatSper1 antibody used in this study is raised against the 1-150 aa region of CatSper1 (blue, Ren et al., 2001). Detection of recombinant FL-CatSper1 and ND-CatSper1 expressed in 293T cells (*right*). (F) A cartoon of the experimental scheme to test NTD truncation of CatSper1. FL-CatSper1 and ND-CatSper1 expressed in 293T cells were solubilized and pulled-down using agarose resin conjugated with HA antibody. The enriched recombinant proteins were incubated with solubilized sperm lysates at 37 °C for 0, 10, 30, and 60 min and subjected to immunoblot. (G) FL-CatSper1 is cleaved at NTD by solubilized sperm lysate. FL-CatSper1 (arrow) decreases while truncated form (arrowhead) increases by incubation with solubilized sperm lysates (*top*). ND-CatSper1 proteins remain largely unchanged under the same conditions (*bottom*). Immunoblotting were performed with HA antibody (E and G). (H) Capacitation-associated CatSper1 degradation is regulated by phosphorylation. CatSper1 degradation is accelerated by PKA inhibition. A PKA inhibitor, H89 (50 μM), enhances capacitation-associated CatSper1 degradation. A protein phophatase1 inhibitor, calyculin A (Caly A, 0.1 μM), prevents the CatSper1 degradation during sperm capacitation *in vitro*. (I-J) Ca^2+^ influx accelerates CatSper1 degradation. (I) CatSper activation during capacitation and Ca^2+^ ionophore treatment (A23187, 10 μM) facilitates the CatSper1 cleavage. (J) Immunodetection of CatSper1 in the spermatozoa incubated under the conditions used in (I). The extent of CatSper1 degradation is heterogeneous in the capacitated sperm cells (*bottom*) compared with A23187-treated uncapacitated sperm cells (middle). (K) Capacitation-associated CatSper1 degradation is blocked by calpain inhibitors (I, II, and III). 20 μM of each calpain inhibitor was treated to sperm during capacitation.

### CatSper1 resides in the subdomains of lipid rafts in mature sperm and processed during capacitation

Sucrose density gradient centrifugation identifies CatSper1 in lipid raft subdomains in mature sperm (Figure 1C). Because cholesterol depletion destabilizes the plasma membrane during sperm capacitation, one simple hypothesis is that the capacitation-associated changes in raft stability and distribution (Nixon et al., 2007) render CatSper1 accessible to a protease activity. Before inducing capacitation, CatSper1 is not processed in sperm cells, probably because the CatSper1-targeting protease activity is normally not in the immediate vicinity to the CatSper nanodomains in the flagellar membrane (Figure 1-figure supplement 1C, E). Supporting this notion, the protease activity readily cleaves CatSper1 by solubilizing the sperm membrane fraction with Triton X-100 (Figure 1-figure supplement 1C).

### The CatSper1 N-terminus undergoes capacitation-associated degradation *in vitro*

We next investigated the location of CatSper1 cleavage and degradation using recombinant CatSper1 proteins and sperm lysates. The CatSper1 antibody used in this study is raised against the first N-terminal 150 amino acids of recombinant CatSper1 (Ren et al., 2001). C-terminal HA-tagged full-length (FL) or N-terminal deleted (ND) recombinant CatSper1 are expressed in HEK 293T cells for pull-down and detection by western blot (Figure 1E, F). Solubilized sperm lysates degrade FL-CatSper1 and result in increased detection of cleaved CatSper1 by HA antibody (Figure 1G; *upper*). In contrast, protein levels of recombinant ND-CatSper1 are not affected by incubation with sperm lysate (Figure 1G; *lower*). These results demonstrate that the cytoplasmic N-terminal domain of CatSper1 is the target region for proteolytic activity in sperm cells. How is the CatSper proteolytic activity regulated?

### CatSper1 degradation involves Ca^2+^ and phosphorylation-dependent protease activity

At the molecular level, capacitation is initiated by HCO_3_^-^ uptake, which activates soluble adenylyl cyclase (sAC), resulting in increased cAMP levels. HCO_3_^-^ also stimulates CatSper-mediated Ca^2+^ entry into sperm cells by raising intracellular pH (Figure 1-figure supplement 1E). We thus examined whether the proteolytic activity requires cAMP/PKA and/or Ca^2+^ signaling pathways. Interestingly, adding a PKA inhibitor H89 or the St-Ht31 peptide, which abolishes PKA anchoring to AKAP during sperm capacitation, accelerated CatSper1 degradation during sperm capacitation (Figure 1H, Figure 1-figure supplement 1D). Consistently, calyculin A, a serine/threonine protein phosphatase inhibitor, suppresses the capacitation-associated CatSper1 degradation. These data suggest that regulation of the proteolytic activity targeting CatSper1 involves protein phosphorylation cascades (Figure 1-figure supplement 1E). Interestingly, adding Ca^2+^ ionophore A23178 to the sperm suspension was sufficient to induce CatSper1 processing even under non-capacitating conditions that do not support changes in PKA activity (Figure 1I, J). Thus, a Ca^2+^ dependent protease that is indirectly regulated by protein phosphorylation such as calpain (Ono et al., 2016) may process CatSper1. We observed that calpain inhibitors prevent CatSper1 from capacitation-associated degradation (Figure 1K).

### CatSper1 degradation correlates with pY development in sperm cells capacitated *in vitro*

The presence and integrity of the CatSper nanodomains, probed by CatSper1 antibody, is an indicator of sperm capability to hyperactivate (Chung et al., 2017; Chung et al., 2014). Inducing sperm capacitation *in vitro* results in a functionally heterogeneous sperm population in which no more than ~15% of cells are hyperactivated (Neill and Olds-Clarke, 1987). This is because individual sperm cells undergo time-dependent changes. The extent to which protein tyrosine phosphorylation (pY) develops and CatSper1 degrades varies with individual sperm cells capacitated *in vitro* (Figure 1J, Figure 2A-C). Notably, we find that sperm cells that maintain intact CatSper1 develop capacitation-associated pY to a lesser degree *in vitro* (Figure 2B, C). This finding is consistent with the reported phenotype of *CatSper1* knockout sperm that exhibit potentiated pY during capacitation (Chung et al., 2014). Thus far, our results suggest that *in vitro* capacitation generates a heterogeneous sperm population in which intact CatSper1 and pY development are inversely correlated in sperm cells at the single cell level. These heterogeneous sperm cells *in vitro* may reflect a collection of the time- and space-dependent changes that sperm undergo in the oviduct (Chang and Suarez, 2012; Demott and Suarez, 1992).

**Figure 2.**
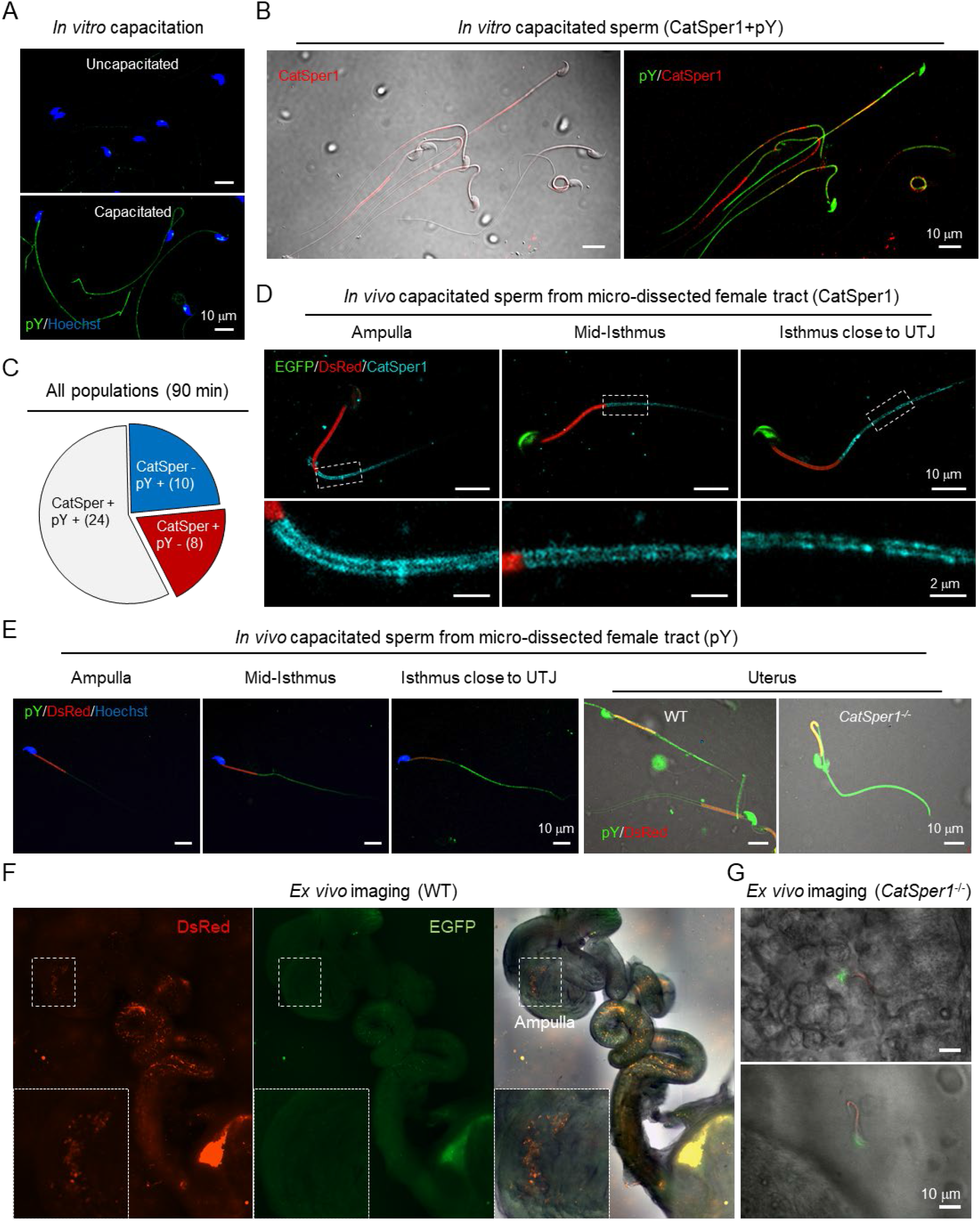
Sperm cells become heterogeneous functionally and molecularly along the female tract. (A) Immunodetection of pY from *in vitro* capacitated sperm. (B) Sperm cells that maintain intact CatSper1 during *in vitro* capacitation exhibit reduced pY development. An image of CatSper1 (red) is merged with the corresponding DIC image *(left*) or pY image *(right*). (C) A pie chart represents expression patterns of CatSper1 and pY in individual sperm capacitated *in vitro*. Sperm number in each group are indicated in parentheses. *Su9-DsRed;Acr-EGFP* WT and *Su9-DsRed;Acr-EGFP CatSper1^-/-^* mice (Chung et al., 2014) were used for mating. Sperm were capacitated *in vitro* for 90 min (A-B). (D-E) Sperm cells capacitated *in vivo* show distinct molecular characteristics along the female tract. The degrees of CatSper1 processing (D) and development of tyrosine phosphorylation (pY) (E) during *in vivo* capacitation were analyzed by immunostaining of the sperm cells at different regions of microdissected female tracts 8h post-coitus. The indicated regions are magnified to show distributions of CatSper1 in sperm cells. Sperm cells that arrived at the ampulla are acrosome reacted and CatSper1 intact (D) and lack pY development (E). Gradual increase of pY is observed in the oviductal sperm located closer to UTJ. Sperm cells that fail to pass UTJ and reside in the uterus show heterogeneous patterns of pY. *CatSper1^-/-^* sperm recovered from the uterus of a mated female show robust elevation of pY. (F-G) WT sperm cells, but not *CatSper1^-/-^* sperm, that arrive at the ampulla are acrosome reacted. *Ex vivo* imaging of female tracts mated with WT (F) and *CatSper1^-/-^* (G) males (8 h post-coitus). (F) WT sperm cells are acrosome-reacted at the ampulla (EGFP-negative, *inset*). (G) A few *CatSper1^-/-^* sperm cells observed at ampulla have intact acrosome (EGFP-positive). Red, DsRed; green, EGFP; Merged, fluorescent images merged with the corresponding DIC image.

### Sperm cells capacitated *in vivo* become heterogeneous along the female tract with distinct molecular characteristics

To assess molecular changes of CatSper1 and pY in the spatially distributed sperm populations along the female reproductive tract, we performed microdissection of the female reproductive tract mated with *Acr-EGFP/Su9-DsRed2* male mice (Hasuwa et al., 2010) 8 hr post coitus and flushed out sperm cells from different regions. By subsequent immunostaining, we found that CatSper1 in the spermatozoa that passed the utero-tubal junction (UTJ) are arranged normally along the tail, mostly protected from degradation, but in decreasing intensity and continuity more towards UTJ (Figure 2D, Figure 1-figure supplement 1E). In striking contrast, pY is not detected in the spermatozoa from the ampulla but appears in the oviductal sperm increasingly towards UTJ (Figure 2E). Absence of EGFP reveals that spermatozoa from the ampulla are fully capacitated and acrosome reacted (AR) but the those in the isthmus are undergoing AR (Figure 2D, F). *Ex vivo* imaging of *Acr-EGFP/Su9-DsRed* sperm in the reproductive tract removed from mated female mice reveals segment-specific patterns of the acrosome status (Figure 2F), consistent with the previous observations that AR initiates in the mid-isthmus (Hino et al., 2016; Muro et al., 2016) and reacted spermatozoa are able to penetrate the zona *in vivo* (Jin et al., 2011). Interestingly, we found that a few *CatSper1^-/-^* sperm cells that managed to arrive at the ampulla are all not acrosome reacted (Figure 2G), supporting the notion that CatSper-mediated Ca^2+^ signaling is required for sperm acrosome reaction (Stival et al., 2018). These results suggest that escape of CatSper1 from the cleavage and subsequent degradation suppresses pY development, enabling sperm to maintain hyperactivation capability, prime AR, and achieve the fertilization *in vivo*.

### 3D *in situ* molecular imaging of gametes in the female reproductive tract

The physiological importance of tracing a small number of spermatozoa progressing to the fertilization site prompted us to seek a method that enables direct molecular assessment of single cells inside the intact female tract. We have adapted tissue clearing technologies to establish three-dimensional (3D) *in situ* molecular imaging systems for fertilization studies (Figure 3, Figure 3 – figure supplement 1, Videos 1-6). We found that various tissue clearing methods (Chung et al., 2013; Murray et al., 2015; Yang et al., 2014) are applicable to the reproductive organs from both male and female mice to preserve gross morphology, and fine cellular and subcellular structures. The cleared tissues preserved protein-based fluorescence and were compatible with labeling with dye and antibodies; growing follicles inside the ovary, oviductal folds and multi-ciliated epithelium, different stages of male germs cells in the seminiferous tubules of the testis and the epididymis are readily detected after clearing and labeling (Figure 3A-C, Figure 3 – figure supplement 1, Videos 1-6). 3D volume imaging of the whole cleared female tract well illustrates the uterine and isthmic mucus and the labyrinths of passages sperm must navigate (Figure 3A, B, Figure 3 – figure supplement 1B, C, Videos 3, 5, 6). Moreover, 3D rendering of the images and digital reconstruction of oviductal surface and central lumen depicts continuous and non-disrupted morphology (Figure 3D, E) consistent with reported dimensions and parameters (Stewart and Behringer, 2012), validating the integrity of the processed oviduct.

**Figure 3.**
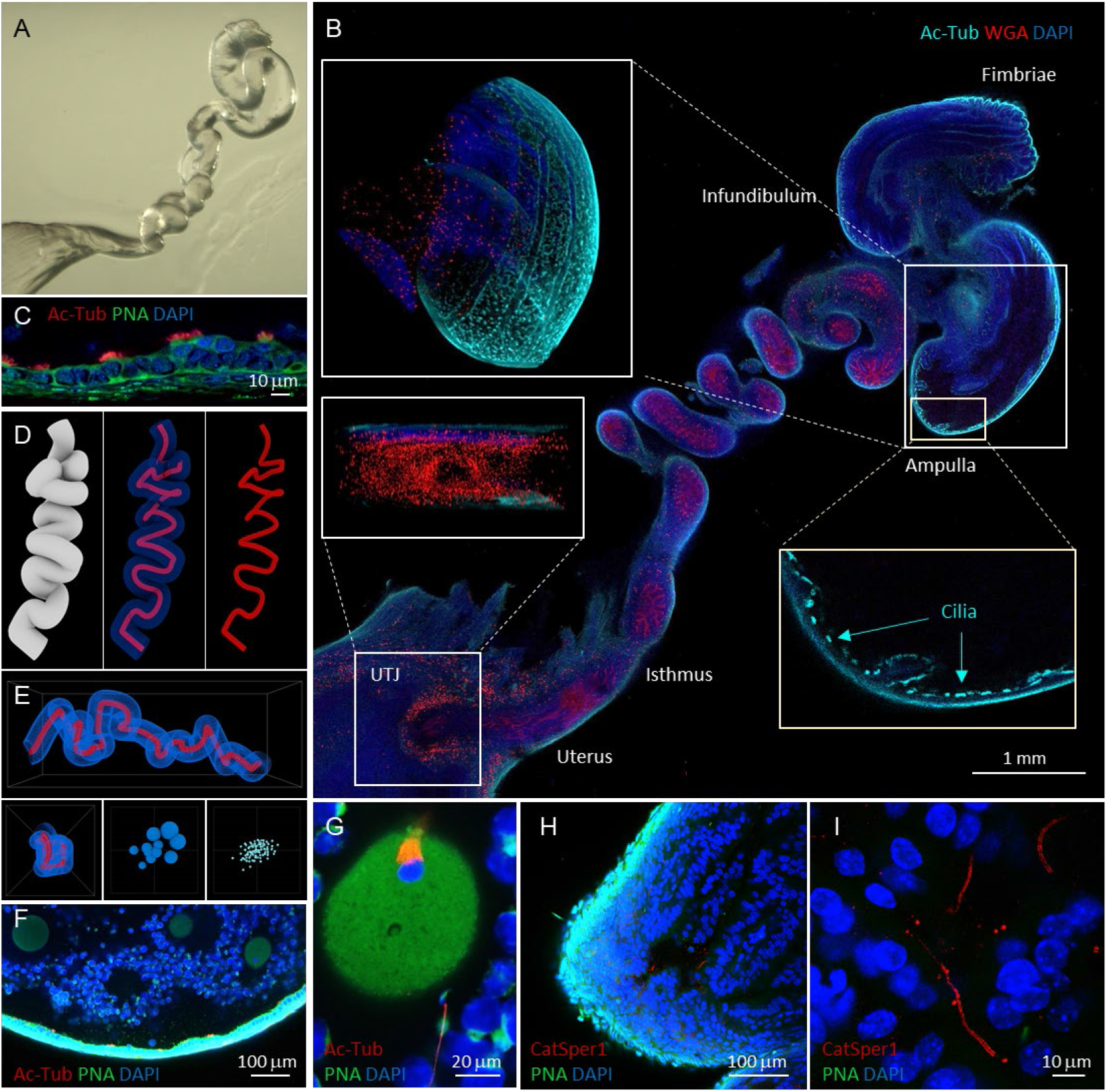
Tissue clearing preserves morphology of female reproductive tract and enables molecular imaging and post-processing of gametes *in situ*. (A) Refractive index-matched cleared mouse female reproductive tract by CLARITY-based tissue clearing. (B) Optical imaging of the cleared female reproductive tract stained by WGA (red), Ac-Tub antibody (cyan) and DAPI (blue), 100x. Insets show cilia stained by Ac-Tub antibody in 2D (*lower right*), a 3D (*upper left*) projection of the ampulla, and a UTJ cross-section (*lower left*). (C) Details of the ampullar epithelium stained by PNA (green), Ac-Tub antibody (red) and DAPI (blue), 400x. (D) 3D digital image reconstruction of the oviduct representing different 3D images rendered for oviductal surface (*left*) and central lumen of oviduct with (*middle*) *or* without (*right*) oviductal volume information. (E) Morphometric and fluorescent signal quantification analysis of the oviduct showing the morphometric meshwork representation of the 3D volumetric data from the oviduct imaging (*upper*), the corresponding side view (*lower left*) and the non-numerical visual representations of the basic volumetric (*lower middle*) and fluorescent (*lower right*) properties. (F) A fluorescent image showing a closer look of the cleared ampulla with oocytes (oocyte magnified in the panel G on the right-most side), 100x. (G) An oocyte with the meiotic spindle; a sperm cell is approaching the *zona pellucida* directly inside the ampulla, PNA (green), anti-AcTub antibody (red) and DAPI (blue), 630x. (H) A tile-scanned confocal image of epithelium of the cleared ampulla (8h post coitus) stained by anti-CatSper1 antibody (red), PNA (green) and DAPI (blue), 100x. (I) Details of the sperm stained directly inside the ampulla by anti-CatSper1 antibody (red). Two linear CatSper domains are clearly recognizable by confocal imaging. Cell nuclei are stained with DAPI (blue); acrosomes are stained with PNA (green). See also Videos 3-7.

We next combined tissue clearing with an *in vivo* sperm migration assay (Chung et al., 2014; Yamaguchi et al., 2009) to molecularly analyze different sperm populations during the fertilization process. Among tested clearing methods, we found that passive clearing of CLARITY-processed reproductive tract from time-mated females retains the location and stability of gametes within the track past UTJ (Figure 3F-I, Videos 4, 7, 8); whole-animal fixation by trans-cardiac perfusion perturbs minimally and rapidly arrests all cellular function while tissue-hydrogel matrix fills the lumen and provides supportive meshwork to prevent gamete loss during subsequent labeling steps (Figure 3 – figure supplement 2). This new *in situ* imaging platform enables capturing a moment of sperm-egg interaction; a spermatozoon that approaches a fertilized egg protruding the 2^nd^ polar body in the ampulla is detected in a cleared female tract 8 h post coitus immunostained by acetylated tubulin antibody (Figure 3F, G, Video 7). CatSper1 antibody specifically recognizes sperm cells transfixed in cleared female tract (Figure 3H). Tissue clearing allows 3D volume imaging of the female tract but does not compromise the resolution. Two linear CatSper1 domains typically observed by confocal imaging are easily observed in the sperm cells inside an ampullar region of the whole cleared female tract (Figure 3I). Thus, the integrity of CatSper1 in sperm cells at different locations along the female tract can be subjected to quantitative analysis.

### Sperm cell that successfully reach the ampulla are CatSper1-intact and acrosome reacted

With this new imaging strategy to detect sperm cells that remain transfixed in the female tract (Figure 3), we investigated acrosome state and CatSper1 integrity in sperm populations directly from the cleared tract of females 8h after mating, focusing on a few anatomically defined regions (Figure 4). Based on the earlier results from micro-dissection or *ex vivo* imaging (Figure 2D-G), we anticipated that sperm cells that successfully reach the ampulla would be CatSper1-intact and acrosome reacted. As expected, most sperm cells located in the ampulla exhibit linearly arranged intact CatSper1 and reacted acrosomes (Figure 4A, *upper*, Video 8). In the middle isthmus, both CatSper1 and acrosome remain intact in most sperm cells, but mixed patterns are observed in some cells (Figure 4A, *middle*, Video 8). Interestingly, acrosome is largely intact in the sperm clusters in the proximal isthmus close to UTJ whereas CatSper1 is barely detected (Figure 4A, *lower*, Video 8). This contrasts with the reduced but readily visible CatSper1 in the sperm from the same region by microdissection (Figure 2D). It is possible that the relatively longer tissue processing time and subsequent labeling could have contributed to lower the signal to noise ratio to a certain degree. Notably, 3D volume imaging of this mid isthmus regions reveals sperm cells aligned in one direction towards the ampulla, providing unprecedented insight into sperm taxis in the fertilization process (Video 8). Our qualitative but semi-quantitative analyses suggest that CatSper1 is largely protected from degradation once in the oviduct; acrosome reaction initiates in the mid-isthmus and is completed in the ampulla before interacting with the oocytes (Figure 4B). These results are consistent with our initial observations from microdissection and *ex vivo* imaging studies (Figure 2D, F), validating the information obtained by our *in situ* molecular imaging platform. Taken together, we conclude that intact CatSper1, lack of pY, and reacted acrosome are molecular and functional signatures of most fertilizing spermatozoa in the physiological context.

**Figure 4.**
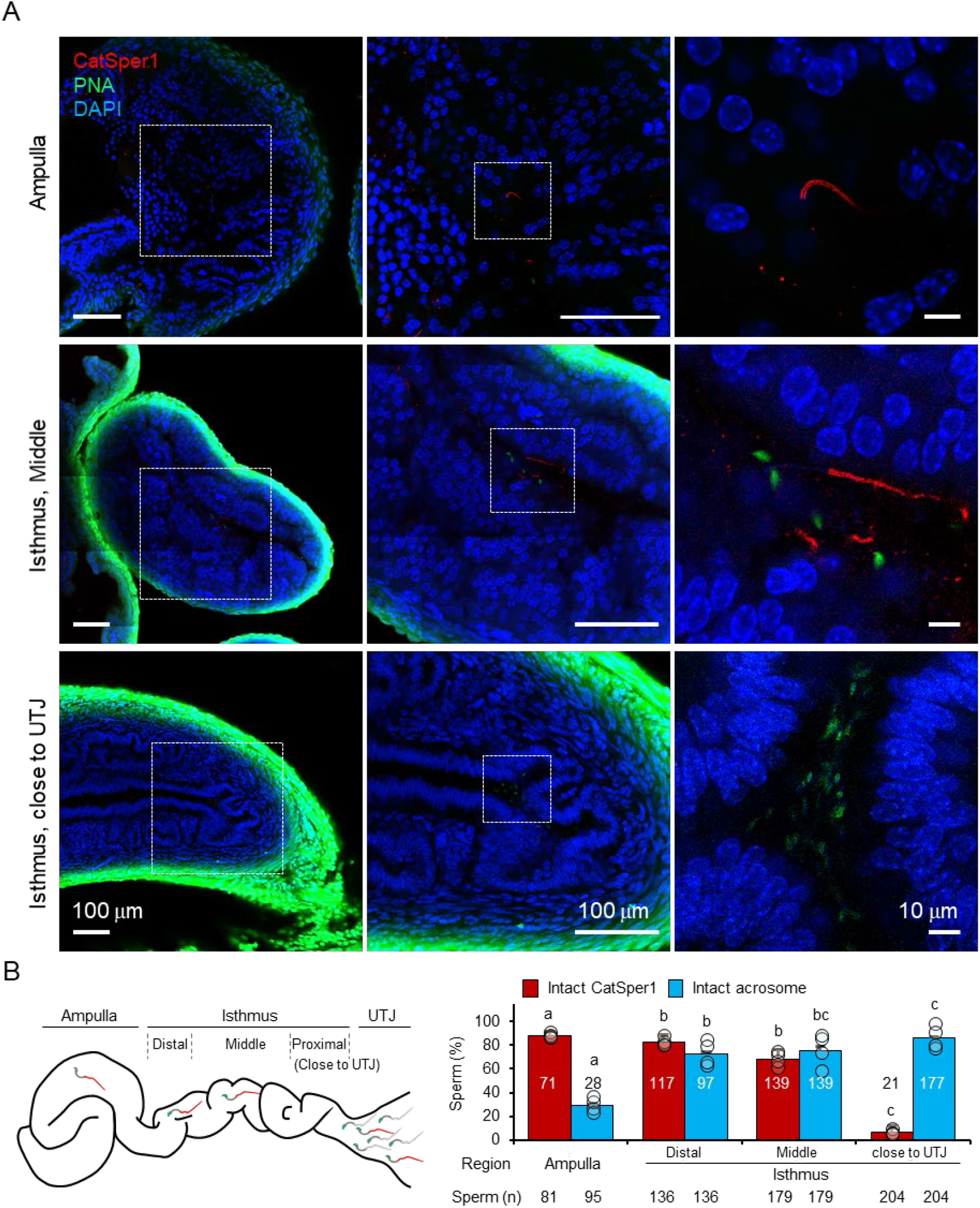
*In situ* molecular imaging of sperm reveals the changes in acrosomal status and CatSper1 fluorescent patterns during capacitation along the female tract. (A) Fluorescent confocal microscope images of acrosome and CatSper1 fluorescent patterns from 3 different regions along the cleared female reproductive tract (Ampulla, *upper*; Middle isthmus, *middle*; Proximal isthmus, *lower*) with different magnifications of the corresponding areas. (B) A cartoon image of the female reproductive tract showing the approximate boundaries between the regions of interest (*left*) used as grouping variable in the subsequent quantification (*right*). The total number of CatSper1-intact sperm (red columns), or acrosome-intact sperm (blue columns), that were counted is shown (bottom). Four independent experiments were performed (n=4). Circles indicate the proportion of sperm cells in each examined site from an independent experiment. Means with different letters indicate significant difference (*P<0.05*) in pairwise comparison between the different regions of female tract. Data is represented to mean ± S.E.M. See also Video 8.

### Automatic detection of sperm in the voluminous female tract using artificial neural network

Processing 3D volumetric fluorescent data presents a significant challenge; analyses of sperm in the female tract includes object identification in the voluminous specimen, object separation from background noise, and object alignment in three dimensions. To address these logistics problems, we took an advantage of the artificial neural network (ANN) approach for automatic localization and signal isolation. We performed a proof-of-principle investigation utilizing CatSper1 distributions in sperm cells from our 3D *in situ* molecular imaging (Figure 5). First, we manually annotated 3D fluorescent signatures of sperm, somatic nuclei and background noise from the original images. These signatures were placed in different abundance models in the ANN 3D training environments (Figure 5A, Figure 5 - figure supplement 1, Video 9) for subsequent ANN training using MatLab ANN module. We performed a supervised iteration process where the sperm locations were predefined in the training environments (Figure 5 - figure supplement 2A). We evaluated the performance of individual ANN according to their sensitivity and specificity in detecting sperm cells and somatic nuclei, and the abundance (voxel occupancy) of noise (Figure 5B, Figure 5 - figure supplement 2B, C). Detection sensitivity is chosen as a major parameter used to evaluate the ANN performance in the training environment simulated with the values similar to those in real samples. The specificity required for sperm detection is lower than the sensitivity, thus provides mainly empty analytical frames that are easily removed manually. After iteration and performance evaluation, we selected the best performing ANN and analyzed images from our experimental samples for which we manually counted sperm number (Figure 5C). The selected ANN is able to recognize all the sperm detected manually and the ANN sensitivity varies around 90% in individual samples (Figure 5-figure supplement 2C), validating the ANN performance. Furthermore, the false-negative detection of sperm all comes from the sperm with dubious signals in the antecedent human eye evaluation; the 90% of the ANN detected sperm expresses well recognizable CatSper1 fluorescent staining patterns (Figure 5C, Figure 5-figure supplement 3A).

**Figure 5.**
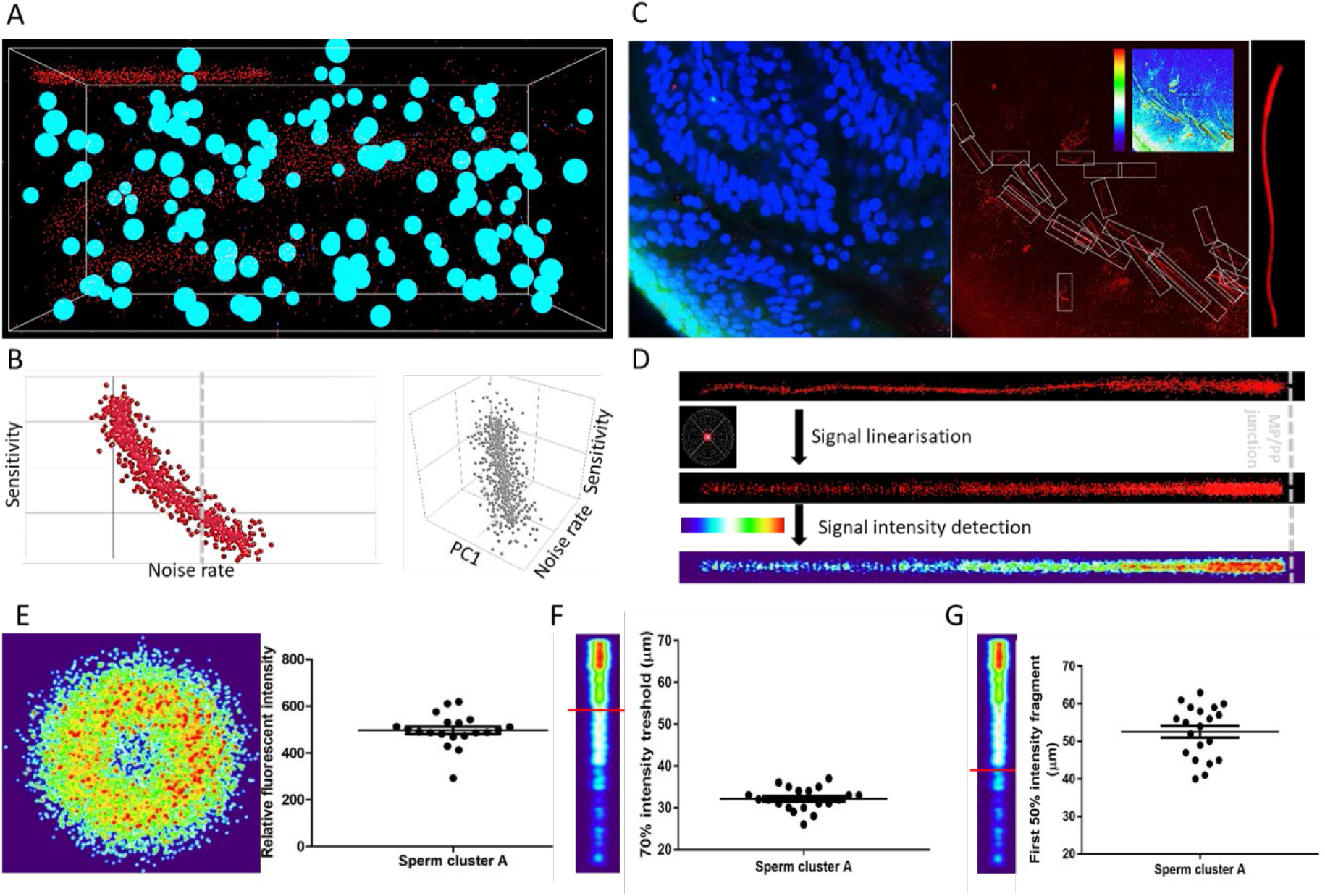
ANN automatically detects fluorescent patterns from 3D volume images of a cleared female reproductive tract, enabling isolation and statistical comparisons of sperm cells. (A) A 3D training environment for ANN emulating sperm cells, somatic nuclei, and noise. (B) Examples of ANN training statistics showing the trend of correlation between the noise rate in the training environment and the sensitivity of ANN (*left*), and a three-dimensional correlation trend between noise rate in the training environment, sensitivity and principal component (PC1) consisting of sperm and nuclei abundancies (*right*). (C) A microscopic focal plane image of the sperm cluster inside the cleared female reproductive tract used for evaluating the ANN performance in real sample (*left*), its superposition in the CatSper1 channel with the individual sperm tails in detection frames with the inset analytical heatmap (*middle*) and the magnification of one of the analytical frames with a CatSper1-positive sperm tail (*right*). (D) Representation of the fluorescent signal in the sperm tail after normalizing individual voxels to signal from the corresponding sperm nucleus (*upper*), after applying linearization and overlay algorithms (*middle*), and heatmap representation of the relative fluorescent intensities among multiple sperm tail (*lower*). (E) Analysis of the relative intensities of the fluorescent signals from sperm located inside the mid-isthmus cleared female reproductive oviduct. The left panel represents the intensity of CatSper1 fluorescent signal in the cross-section of one sperm tail from 20 individual sperm under analysis (middle isthmus). The right panel show the distribution of relative fluorescent intensity of the 20 sperm. (F) Analysis of the continuity of the fluorescent signal along the individual sperm tails; the first fragment of the 70% signal intensity decreases from the midpiece/principal piece interface. (G) The first fragment of the 50% signal intensity decreases from the midpiece/principal piece interface. See also Video 9.

In order to pair each CatSper1 signal containing tail with the head from the same cell in the subsequent analysis, we took the reverse approach to the environment production by removing the detected noise and somatic cell nuclei from the analytical frames (Figure 5-figure supplement 3B). The pre-processed CatSper1 fluorescent signal were then subjected to subsequent alignment, pattern linearization, and intensity detection (Figure 5D, Figure 5-figure supplement 3C). These steps make possible calculation and visual representation of the fluorescent intensity parameters along the sperm tail related to their CatSper1 integrity status (Figure 5E-G).

### ANN-quantified CatSper1 signal reveals a molecular signature of successful sperm *in situ*

The quadrilateral and linear organization of the Ca^2+^ signaling nanodomains discovered by super-resolution imaging of CatSper1 (Figure 6A) is an indicator of a sperm cell’s ability to hyperactivate and fertilize the egg *in vitro* (Chung et al., 2017; Chung et al., 2014). The present study demonstrates that incubating sperm cell under capacitating conditions *in vitro* induces CatSper1 cleavage and degradation, leading to a heterogeneous sperm population (Figures 1, 2). Building on our observations of sperm cells from microdissection, *ex vivo* imaging, and CLARITY-based *in situ* molecular imaging (Figures 2, 3, 4), we hypothesize that CatSper1 is a built-in countdown timer for sperm death and elimination in the female tract; CatSper1 cleavage and degradation, triggered in a time- and space-dependent manner along the female tract, signals to end sperm motility, and ultimately sets sperm lifetime *in vivo*. With our newly developed automated ANN method to obtain high-quality 3D fluorescent images of CatSper1 in the sperm cells from cleared female tract samples, we further tested this idea by quantitatively analyzing the CatSper1 signals *in situ*.

**Figure 6.**
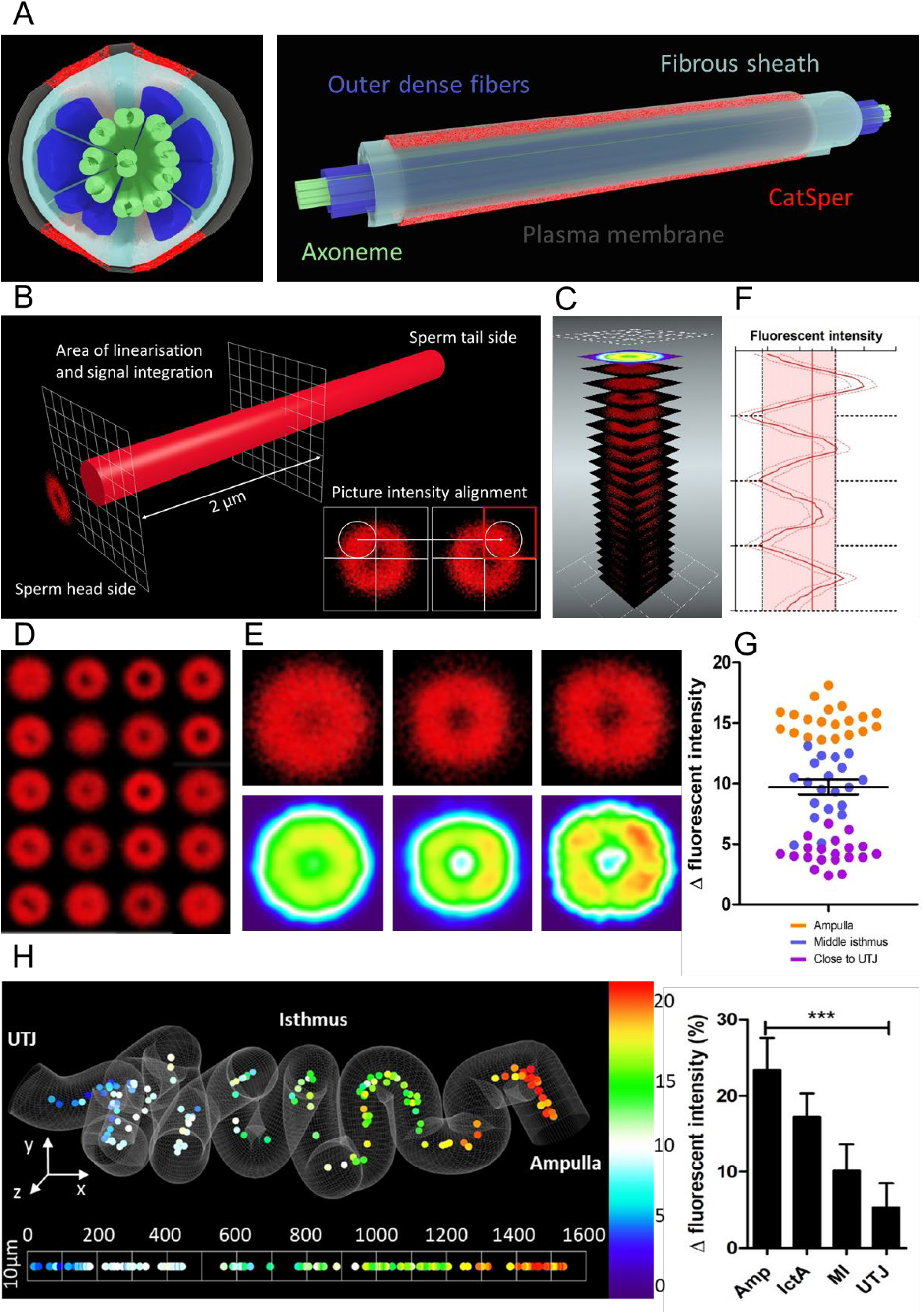
ANN assessment of quadrilateral CatSper nanodomains and Δ fluorescent intensity in sperm population along the cleared female tract conforms to findings by other approaches used in this study. (A) 3D perspective schematic views of quadrilateral CatSper nanodomains. A cross-section (*left*). A side view (*right*). (B) A schematic diagram describing the image processing procedure. (C) An illustration of generating ta heatmap from the pre-processed micrographs. (D) 20 processed micrographs of the CatSper1 signal from the sperm cluster from middle isthmus of the cleared oviduct. (E) Processed micrographs (*upper*) and their corresponding heatmaps (*lower*) from 20 spermatozoa from the oviduct close to UTJ (*left*), middle isthmus (*middle*), and ampulla (*right*). (F) An example of fluorescent intensity analysis of processed images showing the 4 peaks corresponding to four CatSper1 quadrilateral domains and calculated averaged Δ value (transversal red line). (G) Analysis of the fluorescent intensity differences (Δ values; red area in panel F) among 3 sperm populations from ampulla, middle isthmus and isthmus close to UTJ. (H) A topological heatmap showing the integrity of the quadrilateral CatSper domain organization represented by Δ values along the morphometrical space of the cleared oviduct (*left*) with the corresponding inferential statistical analysis of the differences of the signal intensities (Δ values, *right*) among four sperm populations (Amp – Ampulla, IctA – isthmus close to ampulla, MI – middle isthmus, UTJ – utero-tubal junction).

Our *in situ* imaging platform offers the typical resolution that a confocal microscopy can provide; two separated CatSper1 arrangement along the sperm tail (Chung et al., 2017) are detected without any computational processing (Figure 3I, 4A). This encouraged us to develop an analytical procedure to assess the status of CatSper1 quadrilateral and linear distributions. We isolated the fluorescent signal from a proximal region of the principal piece close to the annulus where CatSper1 signal is the most intense (Figure 6B). To superpose the individual cross-sectional images according to the expected 4 intensity peaks, we aligned randomly oriented transversal-projection images by placing the quadrant with the highest fluorescent intensity to upper right corner (Figure 6B, *inset*). The aligned images were then superposed (Figure 6C) and used for statistical purposes to represent quadrilateral arrangement of CatSper1 in individual sperm cells (Figure 6D). The individually processed images of sperm cells from the oviductal regions close to UTJ, middle isthmus, and ampulla, regions were again superposed to create cumulative diagrams and heat maps corresponding to these regions (Figure 6E). They show quadrilateral distribution of enriched CatSper1 signal more clearly from the sperm population in the ampulla compared to the population in the oviduct close to UTJ (Figure 6E, G, H).

To further quantify and statistically analyze our outputs, we divided the pre-processed images of individual sperm cells on 80 round areas (Figure 6 – figure supplement 1A) and calculated fluorescent intensities among them. The quantified intensity from the 80 areas were plotted; the observed 4 peaks (highest intensity) and valleys (lowest intensity) were used to calculate the delta value among them to represent the quality of CatSper1 quadrilateral structure (Figure 6F). Our quantitative analysis (Figure 6G, Figure 6 – figure supplement 1B) shows consistent results with our previous semi-quantitative analysis by manual assignment of the CatSper1 patterns (Figure 4). Together with the whole tissue image processing (Figure 3E), the quantitative analysis clearly visualizes that sperm populations located along the cleared oviduct have statistically different CatSper1 quadrilateral intensity delta values (Figure 6H).

## Discussion

### CatSper1 as a molecular barcode for sperm maturation and transition in the female tract

Testicular spermatozoa undergo maturation and biochemical alterations in the intraluminal environment of the epididymis (Cornwall, 2009). Glycan-modifying enzymes such as glycosidases and glycosyltransferases are present in the epididymal luminal fluid (Tulsiani, 2003). Here we have shown that CatSper1 in mouse sperm is an O-linked glycosylated protein with gradually increasing molecular weight from the testis to the epididymis during male germ-cell development. The different forms of native CatSper1 may represent different degrees of glycosylation. Heterologously expressed CatSper1 cannot reach the plasma membrane, remaining instead at the ER/Golgi (Chung et al., 2017; Chung et al., 2011; Ren et al., 2001). It is intriguing that the molecular weight of recombinant CatSper1 is similar to one of the testicular forms of CatSper1 but bigger than that of the enzymatically deglycosylated and naked polypeptide. O-linked glycosylation takes place in the cis-Golgi for secreted and transmembrane proteins after the protein is folded (Rottger et al., 1998), suggesting that additional modification is required for native CatSper1 to exit the Golgi. Determining the precise identity and modification site may help to clarify the long-sought functional expression of the CatSper channel in heterologous systems. In rodents, sialyltransferase displays maturation-associated quantitative changes (Ram et al., 1989; Scully and Shur, 1988) and sperm lose sialic acid from the surface during capacitation (Ma et al., 2012). Sperm glycoproteins promote sperm migration and survival in the female reproductive tract (Ma et al., 2016). We speculate that mature CatSper1 in sperm contains terminal sialic acid residues, consistent with the small drop in Catsper1 molecular weight during capacitation. The dynamic sugar modifications on CatSper1 may serve as a binding site for decapacitation factors and/or a recognition site during capacitation. Supporting this idea, it was previously shown that mouse sperm lacking CatSper channel cannot pass through the UTJ (Chung et al., 2014; Ho et al., 2009).

Capacitation-associated CatSper1 degradation is blocked by incubation with a 26S proteasome inhibitor, MG-132 (Chung et al., 2014). Now we show that solubilized sperm membrane fraction contains additional proteolytic activities that cleave within CatSper1 NTD. The proteolysis is dependent on Ca^2+^ entry and PKA phosphorylation cascade. A member of calpains, the Ca^2+^ dependent modulatory protease family, may cleave CatSper1, as their proteolytic activity can be positively regulated by PKA (Goll et al., 2003). Among 15 calpain proteins identified in mammals (Ono et al., 2016), calpain1 and calpain11 were previously detected in our sperm proteome (Hwang et al., 2019). Intriguingly, we observed that Ca^2+^ influx by A23187 is sufficient to induce CatSper1 processing under non-capacitating conditions that do not support PKA activation. Increased Ca^2+^ level overrides the phosphorylation effect on calpain1 activity (Du et al., 2018). Calpain11 might be similarly regulated to calpain1, as their domain structures and catalytic residues are conserved (Ono et al., 2016). Since recombinant CatSper1 is cleaved more specifically by sperm lysates, we propose that the testis-specific calpain11 (Ben-Aharon et al., 2006) may target CatSper1.

The effect of CatSper1 truncation on channel activity and sperm motility remains to be determined in future studies. CatSper1 truncation may be coordinated with molecular changes of other CatSper subunits. For example, the protein level of CatSper2, but not CatSper3 or 4, also decreases after capacitation when probed with the antibody recognizing its C-terminal domain (CTD) (Figure 1; Chung et al., 2014). Since the cytoplasmic modulatory subunits, CatSperζ and Efcab9, mainly interact with the channel pore (Hwang et al., 2019), specific processing of the intracellular domains of pore subunits could alter the interactions and subsequent channel activity. Sperm that successfully navigate to the fertilization site in the female reproductive tract and interact with the egg are recognized by intact CatSper1. CatSper1 processing may lead to a loss of control in hyperactivation and eventually end sperm life.

### Physiological function of capacitation-associated tyrosine phosphorylation and acrosome reaction

An increase in pY is one of the various capacitation-associated parameters observed from *in vitro* capacitated sperm cells (Visconti et al., 1995). Subsequently, pY was observed in the flagellum of mouse and human sperm interacting with the oocyte in the medium that supports sperm capacitation and fertilization *in vitro* (Sakkas et al., 2003; Urner et al., 2001). This correlation of pY and the zona binding previously established pY as an indicator of successful sperm capacitation. More recently, however, different observations have been made with *in vivo* approaches. In sows inseminated close to ovulation, spermatozoa found in the UTJ exhibited more phosphorylation in the flagella than those bound to oviductal epithelial cells (OEC), where pY was limited to the equatorial region in the sperm head or no pY was observed (Luno et al., 2013). In mice, the testis-specific tyrosine kinase, Fer, is demonstrated as a master kinase for capacitation-associated pY (Alvau et al., 2016). Surprisingly, homozygous *Fer*-mutant male mice are fertile even though their sperm do not develop pY. All together, these results lead to a new interpretation of the physiological significance of pY: successful sperm capacitation does not require pY development. Determining the precise time and place of pY development in sperm *in situ* would help to elucidate its function in sperm capacitation and fertilization. Here we have shown that sperm, which have capacitated *in vivo* and successfully migrated to the ampulla, are characterized, not only by intact CatSper1, but also by lack of pY development and reacted acrosome. These results coincide with our observations from *in vitro* capacitated sperm cells and other previous studies; pY development inversely correlates with CatSper1 integrity at the single cell level (Figure 2); genetic and pharmacological ablation of Ca^2+^ entry potentiates pY (Chung et al., 2014; Navarrete et al., 2015); AR occurs in mid-isthmus before contacting an oocyte ZP (Hino et al., 2016; Jin et al., 2011; Muro et al., 2016).

Sperm remaining in the female reproductive tract need to be eliminated after fertilization. They may undergo apoptosis and phagocytosis in the female reproductive tract (Aitken and Baker, 2013; Chakraborty and Nelson, 1975) and/or become lost in the peritoneal cavity (Mortimer and Templeton, 1982). pY is reported to mediate apoptosis in immune cells (Yousefi et al., 1994) and cancer cells (Liu et al., 1994). We propose that capacitation-associated global pY development represents degenerating sperm which might concomitantly lose motility. It is intriguing that capacitation-associated reactive oxygen species (ROS) generation activates intrinsic apoptotic cascade and compromises sperm motility (Koppers et al., 2011). Consistent with this idea, ROS inactivates protein tyrosine phosphatase (Tonks, 2005) and enhances pY development in sperm (Aitken et al., 1998). Inhibition of PKA anchoring to AKAPs, which induces CatSper1 truncation and degradation, also suppresses acrosome reaction in capacitating sperm cells *in vitro* (Stival et al., 2018). Thus, CatSper-mediated Ca^2+^ signaling directly or indirectly contributes to sperm acrosome reaction in the female tract. Future work will determine molecular mechanisms by which CatSper channel activity fine-tunes Ca^2+^ signaling to regulate hyperactivated motility, as well as how the Ca^2+^ signaling is linked to coordinate acrosome reaction.

### New *in situ* molecular imaging platform for the study of fertilization and reproduction

Successful development of *in vitro* capacitation and fertilization systems provided fundamental insights into sperm capacitation, fertilization and early embryogenesis. On the other hand, it is evident that the *in vitro* systems have limitations. Sperm numbers required for IVF are much higher than those observed at the fertilization site *in vivo* (Suarez, 2006). Sperm capacitated *in vitro* do not encounter the anatomically and spatially distinct environment of the female reproductive tract, for example, missing their interaction with the oviductal epithelial cells. *In vitro* capacitation also lacks secretory factors from the male and female reproductive tracts that can affect the surface protein dynamics during the capacitation process (Flesch and Gadella, 2000). Mouse models that typically use epidydimal sperm for *in vitro* studies do not contain secretions from male glands. This is in contrast with ejaculated sperm from human and domestic animals. Recent studies have observed sperm behavior in the physiological context through *ex vivo* imaging of sperm in the mouse and bovine oviducts under transillumination (Hino and Yanagimachi, 2019; Ishikawa et al., 2016; Kolle et al., 2009; Muro et al., 2016). Yet this technique is limited in providing molecular information at a single cell level, as live imaging is not easily amenable to direct molecular labeling and 3D volume imaging.

Here, we report new systems to molecularly examine individual sperm cells capacitated *in vivo*. Polymerization of the hydrogel-embedded time-mated female reproductive tract followed by passive clearing provides a stable meshwork to minimally disturb the original location of sperm cells inside the female tract. This approach allowed us to assess the fine organization of CatSper nanodomains in the sperm cells distributed along the female reproductive tract. We showed that both the intensity and the quadrilateral detection of the domains probed by CatSper1 appear as the common pattern of sperm reaching the ampulla and potentially fertilizing the oocyte. The experimental outputs complement the molecular and functional information of sperm released from micro-dissected female tracts and *ex vivo* imaging, identifying molecular and functional signatures of fertilizing sperm in the physiological context. Furthermore, we demonstrate the efficacy of topological heat-map representations of cumulative results by automatic sperm detection and image post-processing and averaging; this method provides statistically robust presentation and interpretation of the volumetric image data.

The present study opens up new horizons to microscopically visualize and analyze molecular events in single sperm cells that achieve fertilization. This will allow us to better understand physiologically relevant cellular signaling pathways directly involved in fertilization. We also have illustrated that the same approach of tissue-clearing based 3D *in situ* molecular imaging is applicable to study gametogenesis *in situ*. Future areas for investigations as natural extensions of the current study are gameto-maternal interaction, development, transport, and implantation of early embryos and maternal-fetal communication. Developing gamete-specific antibodies and/or knockout validated antibodies to probe molecular abundancy and dynamics *in situ* and post-processing tools for various parameters will be critical to this end.

## Materials and Methods

### Animals

*CatSper1-null* (Ren et al., 2001) and *Su9-DsRed;Acr-EGFP* (Hasuwa et al., 2010) mice were generated in the previous study and maintained on a C57BL/6 background. *Su9-DsRed;Acr-EGFP* mice were crossbred with *CatSper1*-null mice to generate *Su9-DsRed;Acr-EGFP CatSper1*-null mice. WT C57BL/6 and B6D2F1 male and CD1 female mice were purchased from Charles River Laboratories (Wilmington, MA) and Jackson laboratory (Bar Harbor, ME). Mice were cared in accordance with guidelines approved by the Yale Animal Care and Use Committees.

### Mammalian Cell Lines

HEK293T and COS-7 cells were purchased from ATCC. They were cultured in DMEM (GIBCO) supplemented with 10% FBS (Thermofisher) and 1x Pen/Strep (GIBCO) at 37 °C, 5% CO2 condition. Cultured cells were used to express recombinant proteins (HEK293T cells) or make total cell lysates (COS-7 cells).

### Antibodies and Reagents

In-house rabbit polyclonal CatSper1 (Ren et al., 2001), CatSper3 (Qi et al., 2007), CatSperε (Chung et al., 2017) antibodies were described previously. Polyclonal CA-IV antibody (M-50) was purchased from Santacruz. Monoclonal antibodies were purchased from BD Biosciences: anti-caveolin1 (clone 2297); EMD Milipore: anti-phosphotyrosine (clon4G10,) anti-acetylated tubulin (clone 6-11B-1), anti-HA agarose (clone HA-7), and Cell Signaling Technology: anti-ubiquitin (clone P4D1), β-actin (clone 13E5), and HA (clone C29F4). HRP-conjugated goat anti-rabbit IgG and goat anti-mouse IgG were from Jackson Immunoresearch. PNA-Alexa 568, WGA-Alexa 555, WGA-Alexa 647, goat anti-mouse IgG (Alexa 488 or 647), and goat anti-rabbit IgG (Alexa 568 or Alexa 647) were from Invitrogen. H89, calyculin A, and Ca^2+^ ionophore A23178 were purchased from Calbiochem. ST-Ht31 was from Promega. Calpain inhibitor II and III were from Enzo life science. All other chemicals were from Sigma-Aldrich unless indicated.

### Epididymal sperm collection and *in vitro* capacitation

Sperm cells were released from caput, corpus, or cauda regions of epididymis in M2 medium (EMD Millipore). To induce capacitation, sperm from caudal epididymis were incubated in human tubular fluid (HTF) medium or M16 (EMD Milipore) containing 25 mM sodium bicarbonate at 37 °C, 5% CO2 condition at 2 x 10^6^ cells/ml concentration for indicated time. Sperm cells were incubated under capacitating conditions with or without the following chemicals: H89 (50 μM), ST-Ht31 (10 μM), Calyculin A (100 nM), calpain inhibitor I (20 μM), calpain inhibitor II (20 μM), or calpain inhibitor III (20 μM). Sperm cells suspended in M2 medium (2 x 10^6^ cells/ml) were incubated with A23187 (10 μM) to induce Ca^2+^ influx under non-capacitating conditions.

### Molecular Cloning

NEB10β bacterial strain (NEB) was used for the molecular cloning. Genomic regions encoding full-length (FL, 1-686 aa) and N-terminal domain deleted (ND, 345-686 aa) mouse CatSper1 were amplified from mouse CatSper1 expression vector (Hwang et al., 2019). The PCR products were subcloned into pcDNA3.1(-) vector using NEBuilder HiFi DNA Assembly (NEB) to express the recombinant proteins tagged with HA at C-terminus (*pcDNA3.1(-)-FL-CatSper1-HA* and *pcDNA3.1(-)-ND-CatSper1-HA*.

### Recombinant protein expression

HEK293T cells were transfected with constructs encoding FL-CatSper1 or ND-CatSper1 to express the recombinant proteins transiently. Polyethyleneimine was used for the transfection following the manufacturer’s instruction as previously.

### Protein Preparation and Western blot

#### Total Protein Extraction

Total proteins were extracted from sperm, testis, and cultured mammalian cells as previously described (Hwang et al., 2019). In short, collected epididymal sperm cells were washed with PBS and lysed in 2X LDS sampling buffer for 10 min at room temperature with agitation (RT). The whole sperm lysates were centrifuged at 14,000 x g for 10 min at 4 °C. Testes were homogenized in 0.32M sucrose and centrifuged at 1,000 x g for 10 min at 4 °C to remove cell debris and nuclei. 1% Triton X-100 in PBS containing protease inhibitor cocktail (cOmplete™, EDTA-free, Roche) was added to the cleared homogenates to make total testis lysate. The lysates were centrifuged at 4 °C, 14,000 x g for 30 min and the supernatant was used for the downstream experiments. Transfected HEK29T cells and COS-7 cells were washed and lysed with 1% Triton X-100 in PBS with protease inhibitor cocktail (Roche) at 4 °C for 1 hr. Cell lysates were centrifuged at 14,000 x g for 30 min. All the solubilized protein lyates from the sources described above were reduced by adding dithiothreitol (DTT) to 50 mM and denature by heating at 75 °C for 5 min (testis and cultured cells) or 10 min (sperm).

#### Discontinuous Sucrose Density Gradient Centrifugation

Discontinuous sucrose density gradient centrifugation was performed as previously described (Kaneto et al., 2008). To isolate and solubilize membrane fraction without using a detergent, cauda epididymal sperm cells washed and suspended in PBS (1.0 x 10^8^ cells/ml) were sonicated 3 times for 1 sec each. Sonicated sperm cells were then centrifuged at 5,000 x g for 10 min at 4 °C and the solubilized membrane fraction in supernatant was collected. The solubilized membrane fraction was pelleted by ultracentrifugation at 100,000 x g for 1 hr at 4 °C and resuspended with PBS. The membrane suspension was mixed with equal volume of 80 % sucrose in PBS. A discontinuous sucrose gradient was layered with the 40%, 30 %, and 5 % sucrose solution from bottom to top in a tube discontinuously. The gradient was ultracentrifuged at 200,000 x g for 20 hr at 2 °C. Proteins collected from each fraction were precipitated with 5 % of trichloroacetic acid, ethanol washed, and dissolved in SDS sampling buffer.

#### Dephosphorylation of Sperm Membrane Proteins

Sperm membrane fractions from 1 x 10^6^ sperm cells prepared as above were treated with protein phosphatase 1, (PP1, 0.1 unit; NEB), protein tyrosine phosphatase (PTP, 5 unit; NEB), or sodium orthovanadate (Na3VO4, 1 mM; NEB) to test dephosphorylation of CatSper1. The membrane fractions were incubated with the phosphatases or Na3VO4 in a reaction buffer containing 20mM HEPES, 0.1 mM EDTA and 0.1mM DTT at 30 °C for the indicated times. The isolated sperm membrane was solubilized by adding Triton X-100 to final 0.1% in PBS (PBS-T) for the indicated times at RT.

#### Enzymatic Deglycosylation

Glycosylation of CatSper1 from cauda sperm was tested using PNGase F (Sigma-Aldrich) and O-glycosidase (NEB). Sperm cells were washed with 1x reaction buffer for each enzyme by centrifugation at 800 x g for 3 min. Sperm pellets were re-suspended with each 1x reaction buffer (20 mM or 50 mM sodium phosphate, pH7.5 for PNGase F and O-glycosidase, respectively) and followed by sonication and centrifugation to collect sperm membrane fraction as described above. Collected supernatants were incubated with denaturation buffer at 100 °C for 5 min to denature glycoproteins before subject to enzymatic deglycosylation. The denatured sperm membrane fractions were incubated with detergent buffer (0.75% IGEPAL CA for PNGase F; 1% NP-40 for O-glycosidase) and each glycosidase at 37 °C for 1 hr. All the enzyme-treated samples were mixed with LDS sampling buffer and denatured after adding DTT to 50 mM at 75 °C for 2 min.

#### In Vitro Proteolysis with Sperm Lysate

Proteolysis of the recombinant CatSper1 protein by sperm lysate were performed as previously described (Chung et al., 2014). Solubilized recombinant FL-CatSper1 and ND-CatSper1 were pulled-down with anti-HA agarose (EMD Millipore) for 1 hr at RT. The enriched recombinant proteins were incubated with 30 μl of sperm lysates solubilized from 3.0 x 10^5^ sperm cells at 37 °C for the indicated times. Sperm lysates were prepared by sonication and incubation in PBS-T without protease inhibitor at 4 °C for 1 hr. After the incubation, mixture of recombinant protein and sperm lysates were mixed to 2X LDS sampling buffer and denatured by adding DTT to 50 mM at 75 °C for 10 min.

#### Western Blot

Denatured protein samples were subjected to SDS-PAGE. Rabbit polyclonal CatSper1 (2 μg/ml), CatSper3 (2 μg/ml), CatSperε (1.6 μg/ml), and CAIV (1:500) antibodies and monoclonal HA (clone C29F4; 1:2,000), caveolin1 (clone 2297, 1:500), acetylated tubulin (clone 6-11B-1; 1:20,000), phosphotyrosine (clone 4G10; 1:1,000), and ubiquitin (clone P4D1; 1:1,000) antibodies were used for western blot. Anti-mouse IgG-HRP (1:10,000) and anti-rabbit IgG-HRP (1:10,000) were used for secondary antibodies.

### Sperm Migration assay

Sperm migration assay was performed as previously described (Chung et al., 2014). Briefly, female mice were introduced to single-caged *Su9-DsRed;Acr-EGFP* males for 30 min and checked for vaginal plug. Whole female reproductive tracts were collected 8 h post-coitus and subjected to *ex vivo* imaging to examine spermatozoa expressing reporter genes in the tract (Eclipse TE2000-U, Nikon).

### Collection of *in vivo* Capacitated Sperm

Female reproductive tracts from timed-mated females to *Su9-DsRed;Acr-EGFP* or *Su9-DsRed;Acr-EGFP CatSper1-null* males were collected 8 h post coitus. Sperm cells were released by micro-dissection of female reproductive tract followed by lumen flushing of each tubal segment (cut into ~ 1-2 mm pieces). Each piece was placed in 50 μl of PBS on glass coverslips and the intraluminal materials were fixed immediately by air-dry followed by 4% PFA in PBS. Ampulla and uterine tissue close to UTJ were placed in 100 μl of PBS and vortexed briefly to release the sperm within the tissues. Fixed sperm cells were subjected to immunostaining.

### Sperm immunocytochemistry

Non-capacitated or *in vitro* capacitated sperm cells on glass coverslips were washed with PBS and fixed with 4% paraformaldehyde (PFA) in PBS at RT for 10 minutes. Fixed samples were permeabilization with PBS-T for 10 min and blocked with 10% normal goat serum in PBS for 1 hr at RT. Blocked sperm cells were stained with primary antibodies, anti-CatSper1 (10 μg/ml) and anti-phosphotyrosine (1:1,000), at 4 °C for overnight, followed by staining with secondary antibodies for 1 hr at RT. Hoechst was used for counterstaining sperm head. Sperm cells were mounted (Vectashield, Vector Laboratories) and imaged with confocal microscopes (Zeiss LSM710 Elyra P1 and Olympus Fluoview 1000).

### Tissue clearing and molecular labeling of the cleared tissues

#### CLARITY

All 3D volume images from the main figures (Figures 3 and 4) were taken from female tracts subjected to CLARITY method (Chung et al., 2013) with slight modification by clearing tissue-hydrogel passively without involving electrophoresis. Timed-mated females (8 h post coitus) and males were subjected to transcardiac perfusion using peristaltic pump. The mice were perfused with each 20 ml of ice-cold PBS followed by freshly prepared hydrogel monomer solution (4% acrylamide, 2% Bis-acrylamide, 0.25% Azo-inhibitor (VA-044, Wako), 4% PFA in PBS). The whole female tract or testis-hydrogel were dissected from animals after perfusion and placed in 10 ml of fresh hydrogel monomer solution for post-fixation. The collected tissues in monomer solution were heated at 37 °C with degassing for 15 min, followed by incubation at 37 °C for 2-3 h for tissue gelation. The gelated tissues were washed with clearing solution containing 200 mM boric acid and 4% sodium dodecyl sulfate (pH8.5) three times for 24 h each by gentle rocking at 55 °C. Cleared tissues were further washed with PBS-T for 24 h. The cleared female tracts were subjected to dye- and/or immunolabeling: the cleared tissues were incubated with CatSper1 (7 μg/ml) or AcTub (1:100) antibodies in PBS-T for overnight at RT, followed by washing with PBS-T for 24 h. Washed samples were stained with the secondary antibodies (1:500) overnight. Fluorescence dye conjugated PNA or WGA were used to detect sugar residues (1:1,000) and DAPI were used for counter staining (1:1000) in PBS-T. Stained tissues were washed and refractive index matched in RIMS solution (Chung et al., 2013) overnight. The index-matched samples were put on imaging chamber filled with RIMS solution and imaged. All cleared tissues were imaged with laser scanning microscope (Zeiss LSM710 Elyra P1). EC plan-Neofluar 10x/0.3, LD LCI Plan-Apochromat 40×1.2, and Plan-Apochromat 63x/1.4 objectives were used for imaging. Tile scanning and z-stacking for volume imaging were carried out with functions incorporated in Zen black 2012 SP2 (Carl Zeiss) and Zen blue 2011 SP1 software (Carl Zeiss) was used for 3D rendering.

#### X-CLARITY

Ovary, testis, and epididymis images (Figure 3 – figure supplement 1A, D-I) were taken from X-CLARITY method, following manufacturer’s instruction (Logos biosystems). Animals transcardially fixed with 4% PFA were post fixed in the fresh fixative for 4-6 h. The post-fixed tissues were then immersed in a modified hydrogel solution (4% acrylamide, 0.25% Azo-inhibitor (VA-044, Wako), 4% PFA in PBS) for 4-6 h. The samples were degassed and polymerized as described in CLARITY method. The gelated tissues were washed with PBS and placed in electrophoretic tissue clearing (ETC) chamber. Tissues in the ETC chamber were cleared by clearing solution described above with active electrophoretic forcing of tissue for 6-8 hr. Cleared tissues were washed and stained with β-actin and WGA.

#### PACT-PRESTO

3D volume images of the oviduct and UTJ (Figure 3 – figure supplement 1B, C) were obtained from the female tract cleared by modified passive ACT-PRESTO (Lee et al., 2016) without involving electrophoretic clearing. In brief, female tracts were fixed in 4% PFA by transcardiac perfusion, followed by post-fixation in fresh fixative solution for 4-6 hours at 4 °C. The post-fixed samples were incubated in the modified hydrogel monomer solution without additional fixative (4% acrylamide, 0.25% Azo-inhibitor in PBS) 4-6 hours at 4 °C. The samples were degassed and polymerized at 37 °C as described in CLARITY method. Hydrogel-infused tissues were cleared with the clearing solution. The cleared tissues were washed with PBS overnight at RT and facilitated labeling is achieved by vacuum-applied negative pressure.

#### SWITCH

3D volume images of testis (Figure 3 – figure supplement 1F) were taken from male mice transcardially perfused and cleared by SWITCH method (Murray et al., 2015). Fixed testes by SWITCH fixative (4% PFA, 1% glutaraldehyde (GA) in PBS) were washed with PBS-T and quenched by 4% glycine and 4% acetamide in PBS at 37 °C for overnight. The quenched samples were passively cleared with SWITCH solution (200 mM SDS, 20 mM Na2SO3, 10 mM NaOH, pH 9.0) 2 times at 60 °C for 3 h each. The cleared tissues were washed with PBS-T for 12-24 h at 37 °C and incubated with refraction index-matching solution (RIMS: 29.4% diatrizoic acid, 23.5% n-methyl-d-glucamine, 32.4% iodixanol). The index matched samples were mounted and imaged.

### Artificial neural network (ANN) image processing

The overall strategy for the artificial neural network (ANN) image processing is described in Figure 4 – figure supplement 2A. The individual signal patterns (sperm, somatic cell nuclei, and noise) were isolated from the original volume images using Zen Blue (Carl Zeiss) and IMARIS software (Oxford instruments) and exported as .obj/.fbx files. The isolated signal patterns were used to generate 3D training environments for ANN by importing different abundancies of the individual components (Figure 4 – Supplementary figure 1) to the 3D environment operating system, Blender 2.79 (https://www.blender.org/); the individual 3D training environment (~10^4^) generated together with the exactly defined coordinates of individual components were exported as .obj/.fbx/Notepad++ files. The ANN training environments were used to develop the ANN detecting the sperm *in situ*. The ANN training was carried out using MATLAB 9.3 (R2017b) software ANN toolbox. The input to the ANN would be virtual *z*-stacks of the produced training environments. The isolated sperm signal patterns were used as as target signature. The supervised training process was performed by comparing the vector coordinates of the individual sperm signatures in the output with the pre-defined vector coordinates of the signatures in the input. This approach also enabled us to evaluate the ANN performance and to quantify signature detection sensitivity and specificity. ANNs with the best performance in detecting the sperm signature were subsequently applied to detect the sperm fluorescent signatures and their post-processing in real volumetric data. In the real environments, selected ANNs showed both sensitivity and specificity around 90%.

## Statistical analyses

Statistical analyses were carried out with one-way analysis of variance (ANOVA) with Tukey post hoc test. Differences were considered significant at p<0.05. For ANN analysis, both parametric (ANOVA; Tukey post hoc) and non-parametric (KW-ANOVA) tests were carried out to evaluate the presented differences; both tests resulted in the same significance output with differences considered significant at p<0.05.

## Acknowledgements

We thank David E. Clapham for sharing reagents and resources, Kwanghun Chung for discussions on tissue clearing methods, Logos Biosystems for use of X-CLARITY system, and Luke L. McGoldrick for critical reading of the draft. This work was supported by start-up funds from Yale University School of Medicine, a Yale Goodman-Gilman Scholar Award-2015, and NIH R01HD096745 to J.-J.C. and by Male Contraceptive Initiative post-doctoral fellowship to J.Y.H.

